# Detecting hidden diversification shifts in models of trait-dependent speciation and extinction

**DOI:** 10.1101/016386

**Authors:** Jeremy M Beaulieu, Brian C O’Meara

**Affiliations:** *National Institute for Biological and Mathematical Synthesis, University of Tennessee*, *Knoxville, TN 37996*, USA; *Department of Ecology and Evolutionary Biology, University of Tennessee*, *Knoxville, TN, 37996-1610,* USA

**Keywords:** hidden states, diversification, comparative methods, speciation, extinction

## Abstract

The distribution of diversity can vary considerably from clade to clade. Attempts to understand these patterns often employ state-dependent speciation and extinction models to determine whether the evolution of a particular novel trait has increased speciation rates and/or decreased their extinction rates. It is still unclear, however, whether these models are uncovering important drivers of diversification, or whether they are simply pointing to more complex patterns involving many unmeasured and co-distributed factors. Here we describe an extension to the popular state-dependent speciation and extinction models that specifically accounts for the presence of unmeasured factors that could impact diversification rates estimated for the states of any observed trait, addressing at least one major criticism of BiSSE methods. Specifically, our model, which we refer to as HiSSE (Hidden-State Speciation and Extinction), assumes that related to each observed state in the model are “hidden” states that exhibit potentially distinct diversification dynamics and transition rates than the observed states in isolation. We also demonstrate how our model can be used as character-independent diversification (CID) models that allow for a complex diversification process that is independent of the evolution of a character. Under rigorous simulation tests and when applied to empirical data, we find that HiSSE performs reasonably well, and can at least detect net diversification rate differences between observed and hidden states and detect when diversification rate differences do not correlate with the observed states. We discuss the remaining issues with state-dependent speciation and extinction models in general, and the important ways in which HiSSE provides a more nuanced understanding of trait-dependent diversification.

---

A key question in biology is, why are some groups are much more diverse than others? Discussions of such questions are often focused on whether there is something unique about exceptionally diverse lineages, such as the presence of some novel trait, which has increased their speciation rate and/or decreased their extinction rates. The BiSSE model (Binary-State Speciation and Extinction; Maddison et al. 2007) was derived specifically as a means of examining the effect that the presence or absence of a single character could have on diversification rates, while also accounting for possible transitions between states. In theory, this model could be used not only for identifying differences in diversification, but also detecting differences in transitions between character states, or even some interplay of the two. In practice, however, it has mainly been used to focus on diversification rate differences (e.g., Goldberg et al. 2010; Wilson et al. 2011; Price et al. 2012; Beaulieu and Donoghue 2013; Weber and Agrawal, 2014).

It is somewhat surprising, perhaps, that studies that employ BiSSE often find that the prediction of a trait leading to higher diversification rates is supported by the data. In fact, all sorts of traits have been implicated as potential drivers of diversity patterns, ranging from the evolution of herbivory in mammals (Price et al. 2012), to the evolution of extra-floral nectaries in flowering plants (Weber and Agrawal 2014), to even the evolution of particular body plans in fungi (i.e., gasteroid vs. nongasteroid forms; Wilson et al. 2011). Some crucial caveats have recently been identified, however. First, Maddison and FitzJohn (2014) raise a statistical concern regarding the inability of these methods to properly account for independence. Consider, for instance, that the carpel, which encloses the seeds of angiosperms, has evolved only once. They argue that the inheritance and pseudoreplication of a single event becomes problematic – even if BiSSE uncovers a significant correlation between the carpel and diversification, it is unclear whether the carpel is an important driver of the immense diversity of flowering plants, or whether this diversity is simply coincidental. It was also pointed out by Beaulieu and Donoghue (2013) that even with many origins of a trait, it could be that only one clade actually has a higher diversification rate associated with the focal trait, which is strong enough to return higher diversification rates for that trait as a whole. In their case, it appeared that plants with an achene (a fruit resembling a bare seed, as in “sunflower seeds”) had a higher diversification rate, but upon subdividing the tree it appeared that this was from the inclusion of one clade in particular, Asterales, which contain the highly diverse Asteraceae, the sunflower family. They argue that it is far more likely that some combination of the achene and another unmeasured, co-distributed trait within Asterales led to a higher diversification rate for achenes as a whole (on this point, also see Maddison et al. 2007). Finally, Rabosky and Goldberg (2015) recently showed that for realistically complex data sets, BiSSE methods almost always find that a neutral trait is correlated with higher diversification. This is, however, largely a consequence of using fairly “trivial” null models that assume equal rates of diversification between character states, which cannot discern the possibility that a complex diversification process is entirely independent from the focal trait.

All these caveats relate to the broader issue of whether the proximate drivers of diversification are really ever just the focal traits themselves. At greater phylogenetic scales this issue seems the most relevant, where the context of a shared trait is unlikely to be consistent across many taxonomically distinct clades. In other words, a character’s effect on diversification will often be contingent on other factors, such as the assembly of particular combinations of characters (e.g., a “synnovation” as defined by Donoghue and Sanderson 2015) and/or movements into new geographic regions (e.g., de Querioz 2002; Moore and Donoghue 2007). Recent generalizations of the BiSSE model (i.e., MuSSE: Multistate Speciation and Extinction; FitzJohn 2012) do allow for additional binary characters to be accounted for when examining the correlation between a binary trait and net diversification. However, it may not always be clear what the exact characters might be, and in the absence of such information, it should be difficult to ever confidently view any one character state as the *true* underlying cause of increased diversification rates.

Here we describe a potential way forward for trait-dependent models of diversification by extending the BiSSE framework to account for the presence of unmeasured factors that could impact diversification rates estimated for the states of any observed trait. Our model, which we refer to as HiSSE (Hidden State Speciation and Extinction), assumes that related to each observed state in the model are “hidden” states that exhibit potentially distinct diversification dynamics and transition rates than the observed states in isolation. In this way, HiSSE is a state-dependent speciation and extinction form of a Hidden Markov model (HMM). As we will show, HiSSE can distinguish higher net diversification rates nested within clades exhibiting a particular character state, can provide more meaningful tests of character-dependent diversification (CID) through its use as a different kind of null model, and thus can provide a much more refined understanding of how particular observed character states may influence the diversification process.

## The Hidden State Speciation and Extinction Model

Despite the important methodological advancement and power afforded by the BiSSE model, it can provide a rather coarse-grained view of trait-based patterns of diversification. Specifically, what may appear like a connection when examining particular characters in isolation may actually be due to other unmeasured factors, or because the analysis included a nested clade that exhibits both the focal character plus “something” else (Maddison et al. 2007; Beaulieu and Donoghue 2013; Beaulieu and O’Meara 2014). This particular point is illustrated in Figure 1. Here the true underlying model is one in which state 0 and 1 have identical diversification rates. However, there is some other trait with states *A* and *B,* and state *B* has twice the diversification rate of *A*. We call this trait a “hidden” trait, because it is not observed in the tip data (though it could be observable if we knew what it was). If state 1 happens to be a prerequisite for evolving state *B* (or even by chance), all the state 0 branches will have state *A*, but some branches in state 1 will have state *A* and some will have state *B*. Thus state 1 actually takes on two states, 1 *A* when the hidden state with higher diversification rate is absent, and 1 *B* when the hidden state with higher diversification rate is present. As indicated by the model, transitions to this unmeasured variable produce nested shifts towards higher rates of diversification *within* clades comprised of species in state 1. Necessarily, BiSSE can only infer parameters for characters that we can observe, and since all origins of state 1 are lumped together, the model infers state 1 as being associated with significantly higher diversification rates. It is associated with higher diversification, but only due to its association with trait B.

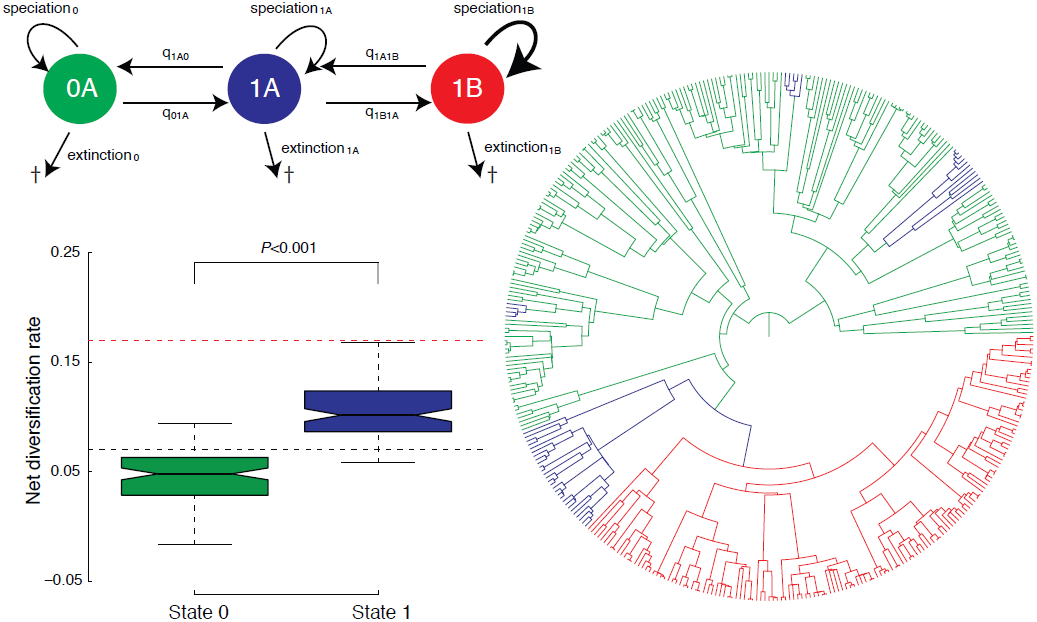

We attempt to solve this problem by deriving an expansion of the BiSSE model that allows for the inference of these hidden states. For the example in Figure 1, we can assume that all observations of state 1 are actually ambiguous for being in each of the possible hidden states, 1*A* (i.e., hidden state absent) or state 1*B* (i.e., hidden state present). We then include transition rates and parameters for the diversification process associated with this hidden state. Our model, which we refer to hereafter as the HiSSE model, is actually a modified form of the MuSSE model (Multi-State Speciation and Extinction; FitzJohn 2012) that extends BiSSE type analyses to allow for multiple binary characters or characters with more than two states. Thus, the HiSSE model can, in theory, have a number of observed states and a number of hidden states (i.e., observed states 0, 1, 2, and hidden states *A*, *B*, *C*, resulting in nine possible state combinations).

Formally, the state space in our model is defined as *o* being the index of the observed state, *o* ∈ 0,1,…, α, and *h* as the index of the hidden state, *h* ∈ *A, B,…,β*. A lineage at time *t* evolves under diversification rates *λ_i_ = λ_oh_* and *μ_i_ = μ_oh_*. Thus, the model has, in general, αβ different diversification rate categories. The likelihood D_N,_*_i_*(*t*) is proportional to the probability that a lineage in state *i* at time *t* before the present (*t*=0) evolved the exact branching structure and character states as observed. Changes in D_N,_*_i_* over time along a branch are thus described by the following ordinary differential equation:

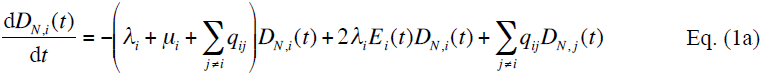
where E*_i_*(*t*) is the probability that a lineage in state *i* at time, goes extinct by the present, and is described by:

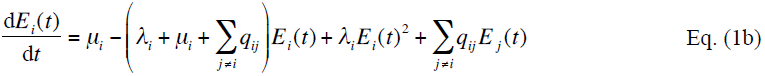

These series of equations are modified from Maddison et al. (2007); they are solved numerically along each edge starting at the tips and moving rootward. The initial conditions for D_N,_*_i_*(0) are set to 1 when the trait is consistent with the observed data, and 0 otherwise. For example, we would set the probability to 1 for both state 1*A* and 1*B* for all species exhibiting state 1 – in other words, the probability of observing a tip demonstrating state 1 is 1 if the true underlying state is 1*A* or 1*B* (note this could be done with states 1*C*, 1*D*, and so forth in this model, although the current implementation in software currently only allows a binary observed character with four or fewer total hidden state combinations). The initial conditions for E*_i_*(0) are all set to zero (i.e., we observe the tip at the present). Incomplete sampling can be allowed by incorporating a state-specific or even clade-specific sampling frequency, *f*, and setting the initial conditions for D_N,_*_i_*(0) as *f_i_* if the corresponding tip, *n,* is in state *i*, and 0 otherwise, and for 1-*f_i_* for E*_i_*(0) (FitzJohn et al., 2009). We assume the sampling frequency of the hidden state to be identical to the state with which it is associated (e.g., *f*_1A_ = *f*_1B_, *f*_0A_ = *f*_0B_).

At each internal node, A, that joins descendant lineages N and M, we multiply the probabilities of both daughter lineages together with the rate of speciation:

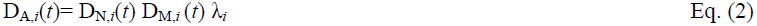
and the resulting values become the initial conditions for the subtending branch. The overall likelihood is the product of D_R,_*_i_*(t) calculated at the root. We condition this likelihood by (1-E*_i_*(t))^2^, which is the probability that the two descendant lineages of the root, evolving under the same estimates of speciation and extinction rates, survived to the present and were sampled (Nee et al. 1994). Finally, we follow the procedure described by FitzJohn et al. (2009) and weight the overall likelihood by the probability that each possible state gave rise to the observed data, although other weights, such as assuming equal probabilities or even “known” fixed probabilities, can also be used. Note that in the absence of a hidden state, our likelihood calculation reduces exactly to the BiSSE model.

We also note that it is rather straightforward to use this framework to implement a SSE version of the “precursor” model described by Marazzi et al. (2012) or the “hidden rates” model (HRM) described by Beaulieu et al. (2013). In the latter case, consider a HiSSE model with “two rate classes”, *A* and *B*. We can define speciation and extinction parameters for states 0*A*, 1*A*, 0*B*, and 1*B*, and then define a set of transitions to account for transitioning between all character state combinations:

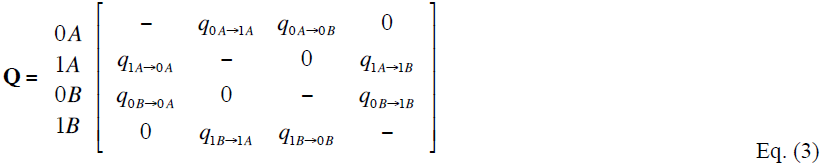

Thus, HiSSE can be used to account for differences in the diversification process while simultaneously identifying different classes of transition rates within a binary character. Our implementation allows for this and more complex models, including those that allow for dual transitions between both the observed trait and the hidden trait (e.g., q_0_*_A_* ↔ q_1_*_B_*).

## Multimodel inference and the issue of appropriate models

An important issue was recently raised by Rabosky and Goldberg (2015), who demonstrated that on empirical trees even traits evolving under a neutral, diversification-independent model will still tend to be best fit by a BiSSE model. FitzJohn (2012) discussed a similar issue of MuSSE falsely assigning diversification rate effects to a character with no such effect. While this behavior is seemingly troubling, it is important to bear in mind that BiSSE, MuSSE, HiSSE, and any other model of state-dependent speciation and extinction are *not* models of trait evolution, but rather joint models for the underlying tree *and* the traits. A trait evolution model like those in Pagel (1994) or Hansen (1997) maximizes the probability of the observed states at the tips, given the tree and model – the tree certainly affects the likelihood of the tip data, but that is the only way it enters the calculation. A trait-based diversification model, on the other hand, maximizes the probability of the observed states at the tips and the observed tree, given the model. If a tree violates a single regime birth-death model due to any number of causes (e.g., mass extinction events, trait-dependent speciation or extinction, maximum carrying capacity, climate change affecting speciation rates, etc.), then even if the tip data are perfectly consistent with a simple transition model, the tip data *plus* the tree are not. In such a case, it should not be surprising that a more complex model will tend to be chosen over a nested simpler model, particularly if the underlying tree is large enough.

Furthermore, as is well known in statistics, rejecting the null model does not imply that the alternative model is true. It simply means that the alternative model fits better. This will often be the case when looking at models in any complex system where the true model may not be one of the included models. For example, Rabosky and Goldberg (2015) showed, among several other empirical examples, that binary characters simulated under a model with each character having no effect on diversification rate on an empirical tree of cetaceans (i.e., whales, dolphins, and relatives) almost always rejected the supposed null model. Though presented as a Type I error (i.e., incorrectly rejecting a true null), it is not. While the chosen character model is wrong, the cetacean tree is almost certainly not evolving with a single speciation and extinction rate for the entire clade (see Slater et al. 2010; Morlon et al. 2011). In other words, BiSSE is correct in saying that the simple model is not correct, but it is very wrong in assigning rate differences to the simulated traits. Nonetheless, even when presented with two bad models, where one assumes constant rates and the other assumes rates are changing rate exactly with character states, BiSSE must still select which model fits the data better than the other.

A simple example may further illustrate this point. Imagine simulating data under a hypothetical model, called “A”, and then comparing the fit of this model against another hypothetical model, called “B”. If model A is a “null model”, but we end up choosing model B, then this would an instance of a “Type I” error – we are incorrectly rejecting the null. Now, imagine simulating data under another distinct model, “C”, but, like before, we only evaluate the fit of model A and model B. Would choosing model B be considered a Type I error? Of course not, because neither model is correct. So clearly the term Type I error (or even Type II error) is outside the region of having any valid meaning in this context.

The reality is that it will be always be a problem comparing any two models and taking the accepted one as the “truth” given a complex underlying reality. Regardless of whether the behavior they observe is properly called Type I error or not, this is the key point of Rabosky and Goldberg (2015). When comparing a simpler model to a more complex model, merely rejecting the simple one does not mean we should accept all the assumptions and interpretations of the alternate one. It is a simple exercise, for example, to simulate a tree using the model of Rabosky (2010) and then fit alternate models such as age-dependent diversification (Alexander et al. 2015) or logistic growth (Rabosky and Lovette 2008) and compare them to a constant rate model; in most simulated trees, the alternate model is chosen (results not shown), even though the processes each assumes were not used to generate the tree.

One potential way forward, at least in regards to the issues Rabosky and Goldberg (2015) pointed out for SSE models, is to provide equally complex models that fit different biological explanations. Take, for instance, the likely complex diversification history of cetaceans. As Rabsosky and Goldberg (2015) demonstrate, when evaluating neutral data simulated on the cetacean tree using a simple, equal diversification rates model, and a complex one with trait-dependent diversification, the data choose the complex one. However, what if there were a complex model that also allowed different diversification rates on the tree, but was completely independent of the diversification process? Then, if the true model were trait-correlated diversification, a model of trait-dependent diversification would be chosen, but if the tree contained some additional complexity, this new trait-independent model might also be chosen some proportion of the time. Again, it bears repeating: *SSE models are not models of trait evolution, but rather, a joint modeling framework for the underlying tree and the trait.* In the case of cetaceans, or really any empirical tree, we do not know the true underlying model that generated the tree, so picking between models of equal rates, trait-dependent, or trait-independent diversification some percentage of the time in no way represents a kind of Type I error. Furthermore, if statistics are done with a focus on parameter estimates rather than model rejection, even if there were some weight for trait-dependent rates, there would also be substantial weight for trait-independent rates as well, and so the average rates across these models should tend to be similar across the possible character states.

Here we propose two character independent diversification (CID) models that are devised to equal the complexity with respect to the number of parameters for the diversification process (i.e., same number of free speciation and extinction rates under the Maddison et al. (2007) parameterization) as a general BiSSE or HiSSE model, but without actually linking them to the observed traits. Thus, these models explicitly assume that the evolution of a binary character is independent of the diversification process without forcing the diversification process to be constant across the entire tree, which is the normal CID used in these types of analyses. The first kind of model, which we refer to as “CID-2”, contains four diversification process parameters that account for trait-dependent diversification solely on the two states of an unobserved, hidden trait (e.g., for speciation rates, λ_0_*_A_* = λ_1_*_A_*, λ_0_*_B_* = λ_1_*_B_*). In this way, CID-2 contains the same amount of complexity in terms of diversification as a BiSSE model. As with BiSSE or HiSSE, these models include the possibility of a variety of constraints for transition rates. These transition rates can be general by allowing them all to be freely estimated, or simplified in various ways such as assuming they are all equal. The second kind of model, which we refer to as “CID-4” contains the same number of diversification parameters as in the general HiSSE model that are linked across four hidden states (e.g., for speciation rates, λ_0_*_A_* = λ_1_*_A_*, λ_0_*_B_* = λ_1_*_B_*, λ_0_*_C_* = λ_1_*_C_*, λ_0_*_D_* = λ_1_*_D_*). The transition rates under this model are set up to account for transitions between the four different rate categories, as well as between the states of the binary character and, in matrix form, they are set up as follows:

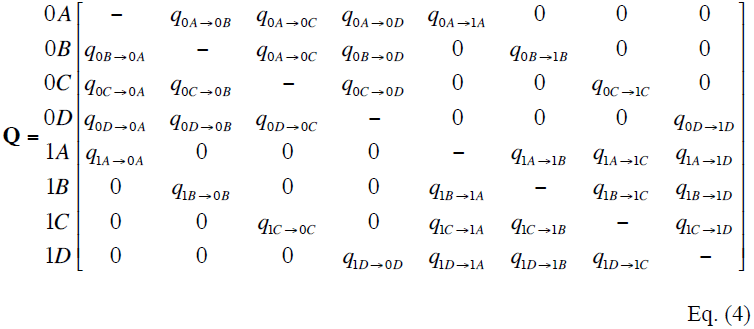

To simplify the number of transitions in the model there are two natural assumptions: assume either all transition rates are equal, or assume there are three distinct transition rates: one rate describing transitions among the different hidden states (i.e., the rates in columns and rows 1-4, and columns and rows 5-8), and two rates for transitions between the observed character states (i.e., one rate for columns 5-8, rows 1-4, and one rate for columns 1-4, and rows 5-8). These or other constraints can be used as part of the CID family of models.

## Implementation

We implemented the above models in the R package *“hisse”* available through CRAN. As input all that *hisse* requires is a phylogeny with branch lengths and a data file that contains the observed states of a binary character. Note that this is an entirely new implementation, not a fork of the existing *diversitree* package, as we employ modified optimization procedures and model configurations. For example, rather than optimizing λ*_i_* and μ*_i_* separately, *hisse* optimizes transformations of these variables: we let τ*_i_* = λ*_i_ +* μ*_i_* define “net turnover”, and we let ε*_i_* = μ*_i_ +* λ*_i_* define the extinction fraction. This reparameterization alleviates problems associated with over-fitting when λ*_i_* and μ*_i_* are highly correlated, but both matter in explaining the diversity pattern (e.g., Goldberg et al. 2010; Beaulieu and Donoghue 2013). With empirical data we often see good estimates for diversification rate but correlations for birth and death rate estimates. For example, Beaulieu and Donoghue (2013) (their Figure S2) showed that the confidence region for birth and death rates tightly follows a diagonal line, with different characters having lines of the same slope but different intercepts. This leads to a behavior where looking at the confidence or credibility intervals for birth or death show overlap between the characters but intervals for diversification rates do not overlap between characters. However, the only way to fit this in the common parameterization is with two birth rates and two death rates. Reparameterizing, as we do, allows us to have the same turnover rate for both states but estimate different diversification rates, resulting in a less complex, but better fitting, model. Thus, users specify configurations of models that variously fix τ*_i_* and ε*_i_.* Note that estimates of τ*_i_* and ε*_i_*, can be easily backtransformed to reflect estimates of λ*_i_* and μ*_i_* by

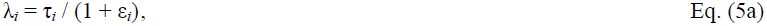

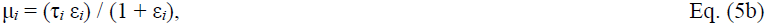
which can be used to derive other measures of diversification process parameters, such as net diversification (λ*_i_*-μ*_i_*). One of the *diversitree* authors (Sally Otto, personal communication) noted that Hardy and Otto (2014) tested a hidden model that only affected transition rates within the MuSSE framework. Note that we have not directly compared the two implementations, though we have confirmed that MuSSE and HiSSE return the same likelihood when all states are observed.

The *hisse* package assigns the probability of each ancestral state to an internal node using marginal ancestral state reconstruction (Yang et al. 1995; Schluter et al., 1997) under the chosen diversification with trait evolution model. These probabilities can be used to “paint” areas of a phylogeny where the diversification rate has increased or decreased due to some unmeasured character. In our implementation the marginal probability of state *i* for a focal node is the overall likelihood of the tree and data when the state of the focal node is fixed in state *i*. Note that the likeliest tip states can also be estimated: we observe state 1, but the underlying state could either be state 1*A* or 1*B*. Thus, for any given node or tip we traverse the entire tree as many times as there are states in the model. As the size of the tree grows, however, these repeated tree traversals can slow the calculation down considerably. For this reason, we also include a function that allows the marginal calculation to be conducted in parallel across any number of independent computer processors.

## Simulations

We evaluated the performance of the HiSSE model by simulating trees and character states under various scenarios and then estimating the fit and bias of the inferred rates from these trees. Our initial simulations first tested scenarios of the single hidden state situation described in Figure 1; the known parameters for each scenario are described in detail in Table 1. We included BiSSE scenarios to test whether we could correctly conclude that there was no support for a HiSSE model in the absence of a hidden state in the generating model. For each of these scenarios, trees and trait data were simulated using *diversitree* (FitzJohn 2012) to contain 50, 100, 200, or 400 species, with 100 replicates for each taxon set. When the generating model included a hidden state, we simulated trees that could transition between three possible states: 0, 1, or 2. After each simulation replicate was completed, we created the hidden state by simply switching the state of all tips observed in state 2 to be in state 1.

**Table 1.**
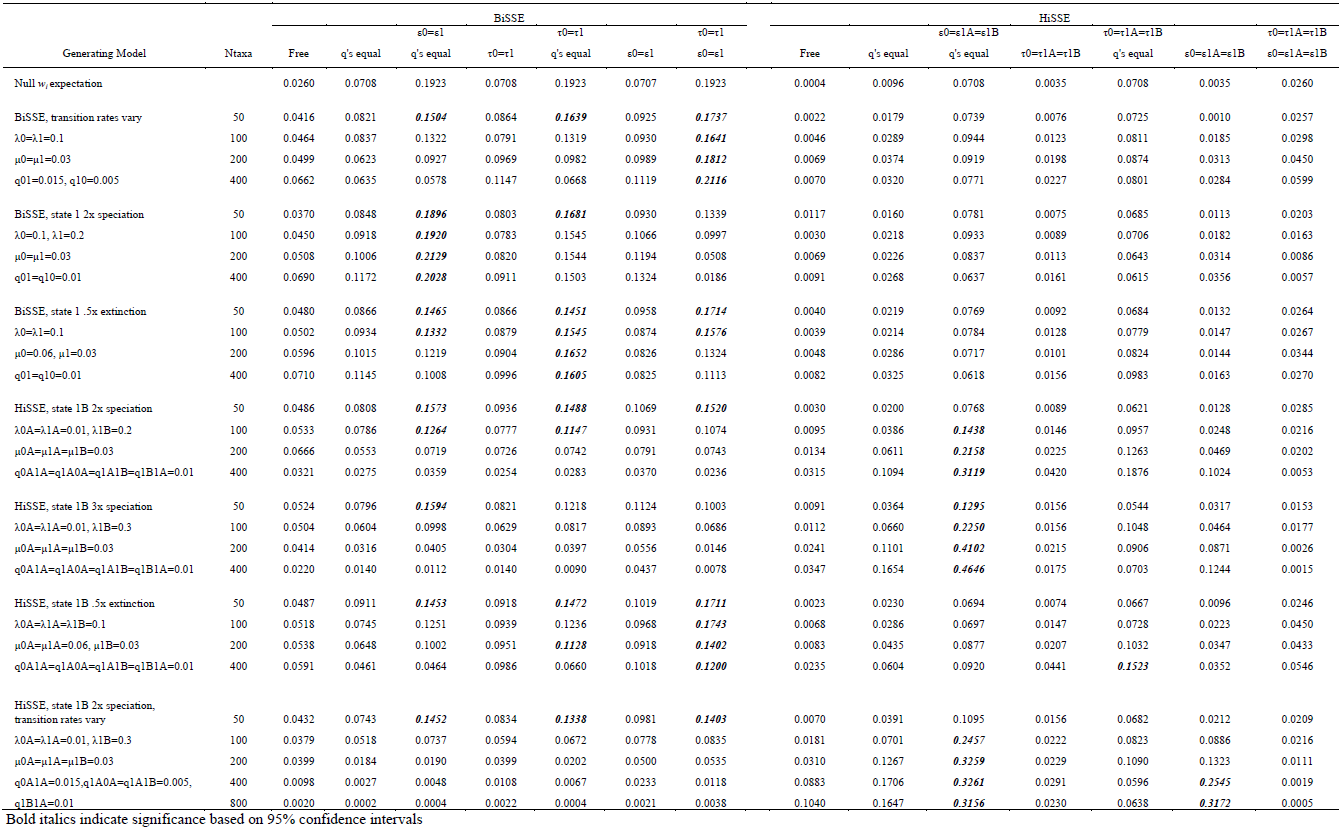
Summary of model support for simulated scenarios with and without a hidden trait, where calculating the average Akaike weight (*w_i_*) for all models assessed the fit. We also calculated a null expectation of the Akaike weight as the average Akaike weight if we assumed an equal likelihood across all models. Thus, the null expectation is based solely on the penalty term in the AIC calculation.

Each simulated data set was evaluated under the generating model, as well as 13 additional models that variously added, removed, or constrained certain parameters. The entire model set is described in Table 1. For all models under a given scenario, model fit was assessed by calculating the average Akaike weight (*w_i_*), which represents the relative likelihood that model *i* is the best model given a set of models (Burnham and Anderson 2002). We also calculated a null expectation of the Akaike weight across our model set, as the average Akaike weight assuming an equal likelihood across all models. Thus, our null expectation is based solely on the penalty term in the AIC calculation – in the absence of information from the model, we would expect to see these weights returned, rather than equal weights for all models. Moreover, since BiSSE is nested within HiSSE, they could return the same likelihood, so even with infinite amounts of data the weight of the HiSSE model should drop to this null expectation, but, counterintuitively, not to zero.

We also conducted a second set of simulations that specifically evaluated the performance of the general HiSSE model and our two CID models of CID diversification. Our goal was to test how much model weight the trait-independent models exhibited under scenarios of trait-dependent diversification. We were concerned at the outset that our CID models would always fit at least as well, if not better, than a trait-dependent BiSSE or HiSSE model. That is to say, these models were constructed such that they are not constrained by character states and can assign rates wherever they want in order to maximize the likelihood. Table 2 describes the known parameters for the three scenarios evaluated. Again we relied on *diversitree* to simulate trees and trait data that contained 400 species, repeated 100 times, with transitions allowed between four possible states, 0, 1, 2, or 3. After each simulation replicate was completed, we created the hidden state by simply switching the state of all tips observed in state 2 to be in state 0, and all tips in state 3 to be in state 1.

**Table 2.** Summary of the model support for simulated scenarios that tested the performance of the general HiSSE model and two CID models (CID-2, CID-2; see text). All data sets tested contained 400 species, and calculating the average Akaike weight (*w_i_*) for all models assessed the fit. As with Table 1, we calculated a null expectation of the Akaike weight as the average Akaike weight if we assumed an equal likelihood across all models.

| Generating Model | Equal rates | BiSSE |  | CID-2 |  | CID-4 |  | HiSSE |  |  |  |
| --- | --- | --- | --- | --- | --- | --- | --- | --- | --- | --- | --- |
| | $\tau_0=\tau_1$ | Free | $\varepsilon_0=\varepsilon_1$ | $\tau_A, \tau_B$ | $\tau_A, \tau_B$ | $\tau_A, \tau_B, \tau_C, \tau_D$ | $\tau_A, \tau_B, \tau_C, \tau_D$ | Free | $\varepsilon_0A=\varepsilon_1A=\varepsilon_0B=\varepsilon_1B$ | $\tau_0A=\tau_1A=\tau_0B$ | $\tau_0A=\tau_1A=\tau_0B$ |
| | $\varepsilon_0=\varepsilon_1$ | | | $\varepsilon_A, \varepsilon_B$ | $\varepsilon_A=\varepsilon_B$ | $\varepsilon_A, \varepsilon_B, \varepsilon_C, \varepsilon_D$ | $\varepsilon_A=\varepsilon_B=\varepsilon_C=\varepsilon_D$ | | | $\varepsilon_0A=\varepsilon_1A=\varepsilon_0B$ | $\varepsilon_0A=\varepsilon_1A=\varepsilon_0B=\varepsilon_1B$ |
| Null $w_i$ expectation | 0.1426 | 0.0865 | 0.1426 | 0.0865 | 0.1426 | 0.0193 | 0.0865 | 0.0117 | 0.0525 | 0.0865 | 0.1426 |
| BiSSE, state 1 2x speciation<br>$\lambda_0=0.1, \lambda_1=0.2$<br>$\mu_0=\mu_1=0.03$<br>All q's = 0.01 | 0.0085 | 0.1995 | **0.3313** | 0.0046 | 0.0089 | 0.0001 | 0.0012 | 0.0255 | 0.0917 | 0.1298 | 0.1989 |
| CID-2, state B 2x speciation<br>$\lambda_0A=\lambda_1A=0.1, \lambda_0B=\lambda_1B=0.2$<br>$\mu_0A=\mu_1A=\mu_0B=\mu_1B=0.03$<br>All q's = 0.01 | 0.0505 | 0.0445 | 0.0568 | 0.1836 | **0.3107** | 0.0063 | 0.0656 | 0.0322 | 0.1461 | 0.0407 | 0.0631 |
| HiSSE, state 1B 2x speciation<br>$\lambda_0A=\lambda_1A=\lambda_0B=0.1, \lambda_1B=0.2$<br>$\mu_0A=\mu_1A=\mu_0B=\mu_1B=0.03$<br>All q's = 0.01 | 0.0049 | 0.0957 | 0.1327 | 0.0073 | 0.0135 | 0.0002 | 0.0032 | 0.0417 | 0.1199 | 0.2265 | **0.3544** |
| "Worst-case" trait-independence<br>Initial $\lambda=0.1$, drawn from lognormal<br>$\varepsilon=0.3, q_01=q_{10}=0.01$ | 0.0005 | 0.0003 | 0.0007 | 0.0897 | 0.1562 | 0.0544 | **0.3996** | 0.0563 | 0.2213 | 0.0067 | 0.0142 |

Bold italics indicate significance based on 95% confidence intervals that bracket the mean Akaike weight.

We also included a scenario that was designed to test whether the general HiSSE model is immune to empirical issues of spurious assignment of importance to state combinations that have no actual effect on diversification. In other words, is HiSSE still favored in situations where the trees and traits evolved under a very different model than the one used for inference? Here we generated trees containing 400 taxa using code from Rabosky (2010). A symmetric Markov model for trait evolution alone (no influence by diversification) was used to simulate binary traits on this tree (using the R package *geiger;* Harmon et al. 2008; Pennell et al. 2014). The Rabosky (2010) model was used, as it is very different from the model assumed by BiSSE/HiSSE; speciation rates evolve gradually on branches, rather than moving discretely between distinct levels based on a trait (hidden or not). Though the rate change is gradual under the Rabosky (2010) model, the speciation rate does not evolve under a Brownian, Ornstein-Uhlenbeck, or similar processes, but in a heterogeneous way that depends on the timing of speciation events (Beaulieu and O’Meara 2015). Thus, as with many empirical data sets, this diversification model is quite different from HiSSE’s model, providing a difficult challenge. The Rabosky (2010) model has also been very influential in affecting biologists’ attitudes towards estimating extinction rates, and so we include it as a semi-realistic “worst-case” scenario. Each simulated data set was evaluated under multiple models that variously added, removed, or constrained certain parameters, with model fit again being assessed by calculating the average Akaike weight (*w_i_*). The entire model set is described in Table 2. For simplicity, all models in this set assumed equal transition rates.

In regards to the parameter estimates in all our simulations, comparisons between the model-average of the parameters against the known parameters provided an assessment of the bias and precision of the inferred parameters. However, rather than averaging parameters across the entire model set, we only averaged across models that included similar parameters. For example, when estimating the bias in the HiSSE scenarios we only model-averaged parameters for models that included the hidden state. This required reformulating the Akaike weights to reflect the truncated model set. Finally, we also assessed the reliability of the ancestral state reconstructions by comparing the true node states from each simulated tree to the marginal probabilities calculated from the model-averaged parameter estimates.

## Assessing model fit

Our main focus is estimating parameters well, but this is aided by picking the true model with high weight when it is in the set (though, of course, for real data the true model is always more complex than any examined one). From a model comparison perspective, our first set of simulations indicated that data sets that lack hidden states could generally be distinguished from those that do, especially with larger data sets (Table 1). When the generating model is a BiSSE model with only two observable states, there were low levels of support for all seven HiSSE models. In fact, as sample size increased, the average Akaike weight of the HiSSE models converged towards the null Akaike weight based only on the penalty term (Table 1). When evaluating data sets that included a hidden state, the ability to correctly favor a HiSSE model over any of the BiSSE models depended not just on the size of the data set, but also the underlying generating model. For example, when the generating model assumed a hidden state with higher speciation rates, data sets that contained 200 or more taxa were required to provide strong evidence for a HiSSE model that varied the turnover rate (Table 1). However, when the main effect of the generating model assumed lower extinction fractions for the hidden state there remained strong support for a BiSSE model that assumed both equal turnover rates and extinction fractions. Interestingly, when we simulated under a HiSSE scenario that combines the processes of higher speciation rates and asymmetrical transition rates, the issue of incorrectly favoring a BiSSE model disappears (Table 1).

We also found that HiSSE was able to distinguish between generating models that assumed diversification rates differences are trait-dependent (e.g., when speciation is two-fold greater for state 1*B*), or when the diversification rate difference is trait-independent and due to simply to the presence of a hidden trait only (e.g., when speciation is two-fold greater for hidden state B) (Table 2; Table S1). And, contrary to our concerns, when the generating model assumed some form of trait-dependent diversification neither the CID-2 nor the CID-4 model had much support.

For the “worst-case” scenario, which simulated a neutral binary character along trees generated from a complex heterogeneous rate branching process, it clearly falls into the zone where BiSSE has unwelcome behavior (94% of data sets favored a BiSSE model over a single-rate diversification model, and the average Akaike weight for state-dependent models across data sets was 92.0%; see Table S1). Thus, these simulation conditions create difficult data sets of the sort used by Rabosky and Goldberg (2015). The addition of merely the CID-2 model dramatically improved performance, where the average weight for the BiSSE models went from 92.0% to 1.3%. When the full HiSSE model set was examined, we found that the CID-4 trait independent model had the highest support on average across the entire set of models (Table 2; Table S1). However, if we combine support for both this CID model and the CID-2 model, they cumulatively account for 70% of the total model weight. Both of these models are independent of any particular character state, but assume shifts among distinct levels, and therefore more closely resemble the conditions by which these data sets were generated.

We note, however, that there was some support for a trait-dependent HiSSE model with the same number of diversification parameters (Table 2; Table S1). And, in spite of the Akaike weight across all data sets showing greater support for the CID-4 model on average, 29% of the individual data sets would favor some form of the general HiSSE model (Table S1). However, there was only a 1.02% difference in the mean diversification rate between states 0 and 1 (though, focusing on just the set of trees for which a HiSSE model was best, the percent difference is 16% – still relatively small for diversification studies, but clearly not as good). Taken together, these results indicate that in these “worst-case” trait-independent scenarios, likely encountered in many empirical data sets, the inclusion of the CID models can attenuate the issue of spuriously accepting trait-dependent diversification. The addition of models that allow some dependence with the observed traits and some with hidden traits, the rates of falsely assessing some state-dependence somewhat increases, but with the benefit of being able to better detect hidden effects when they are true (see earlier simulation results).

## Estimating model parameters

It is important to point out that in all simulation scenarios we never recovered strong support for a model consistent with the model that actually generated the data. That is, whether we vary the speciation rate or the extinction rate, which affects both net turnover and extinction fraction, we always found support for a simplified model that either allowed τ to vary or ε to vary, but not both. This is likely a consequence of the uncertainty and upward biases in estimates of extinction fraction (Fig. 2-4), making it difficult for the model to infer multiple extinction fractions, at least with smaller data sets.

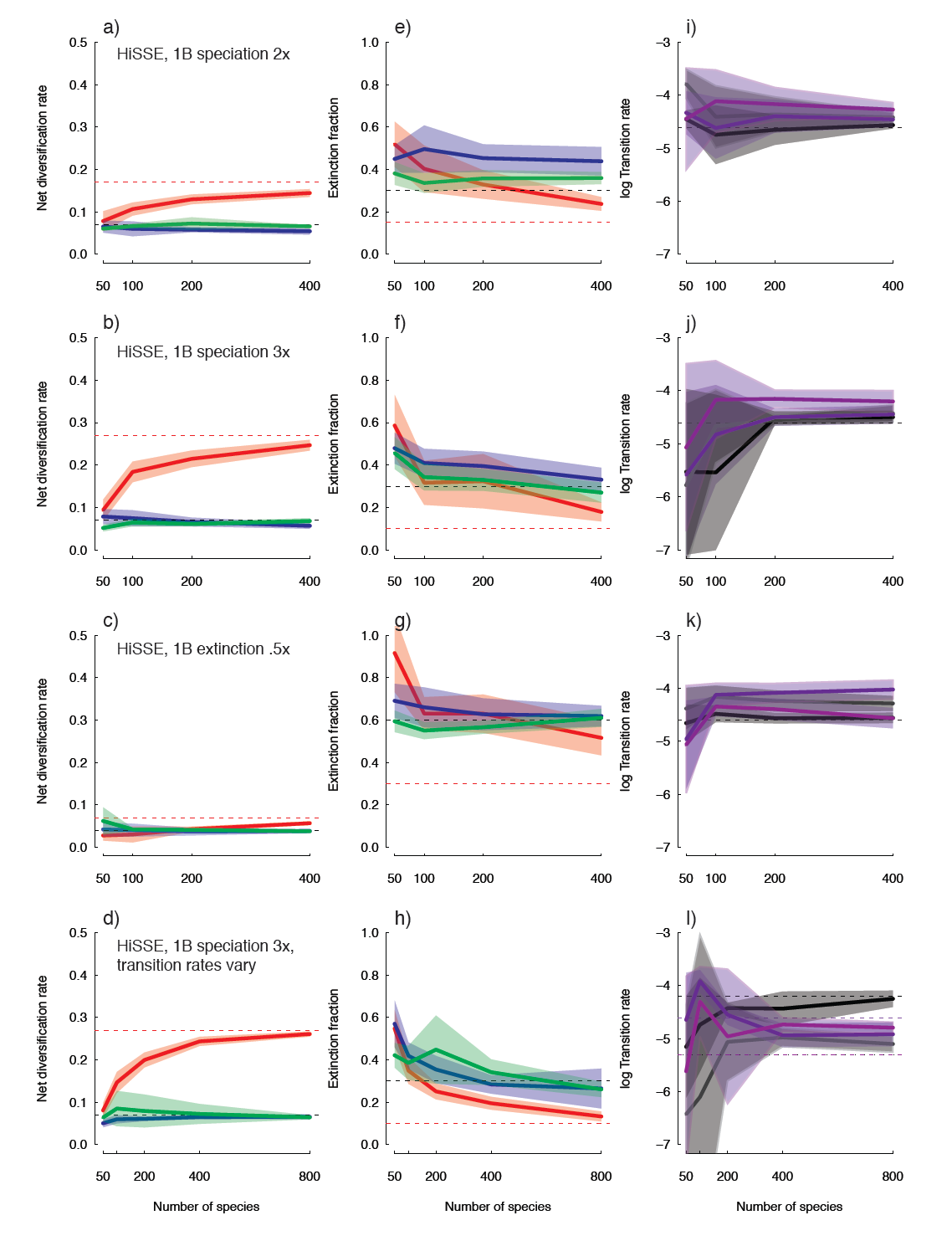

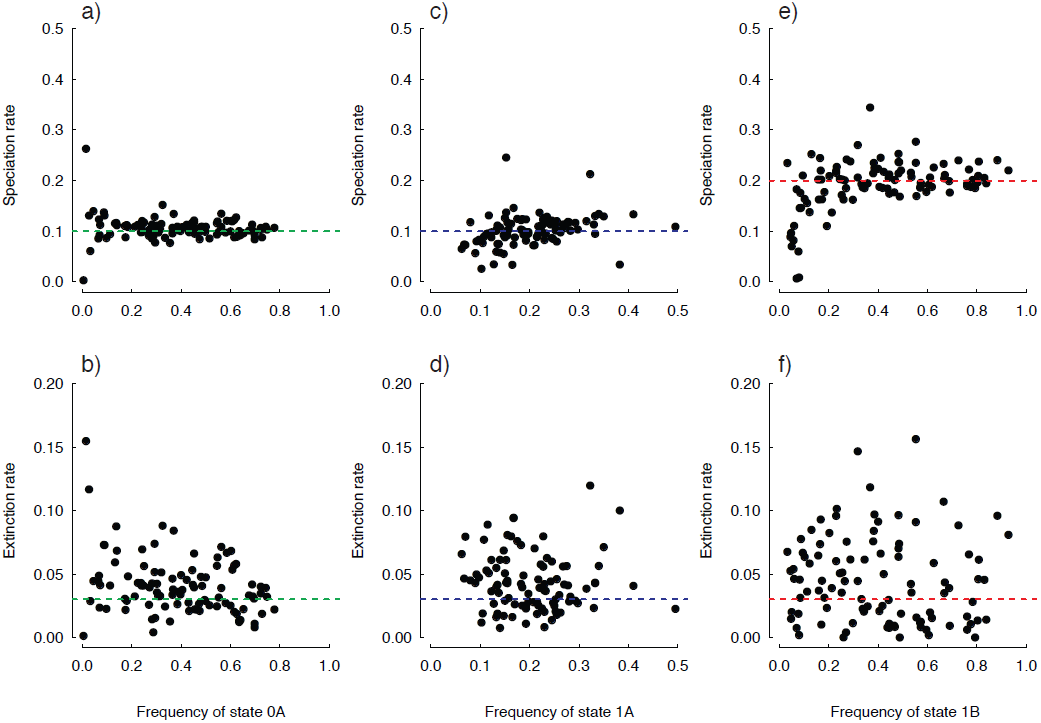

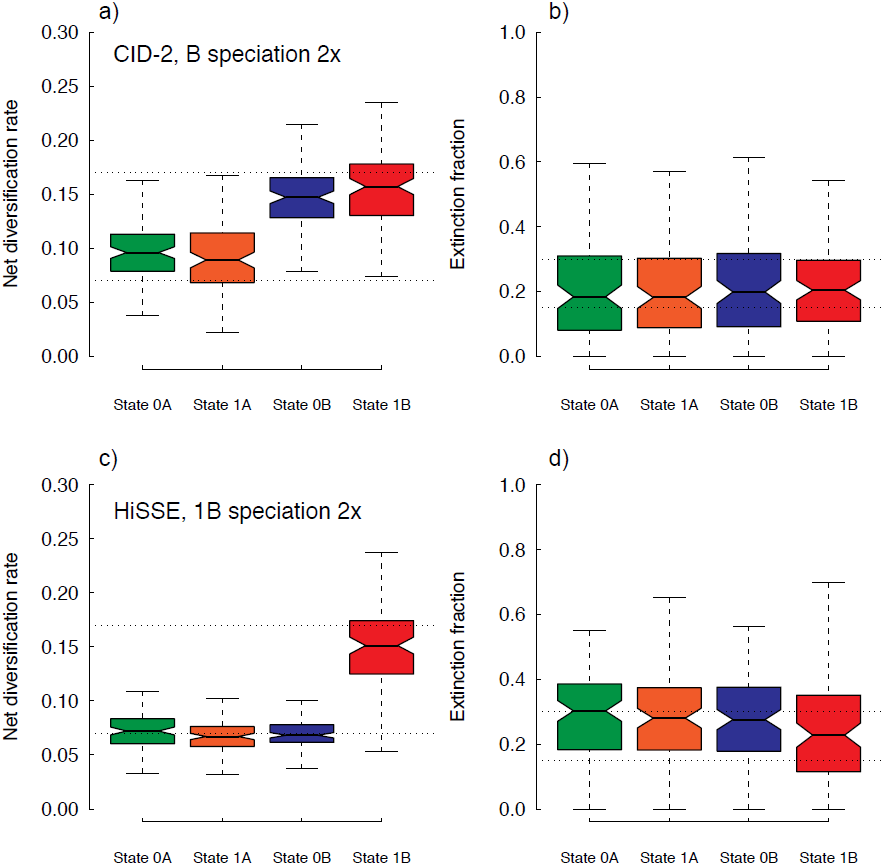

It is well known that there can be difficulties, generally, in obtaining precise estimates of the extinction rates (i.e., μ*_i_*) under the BiSSE framework (Maddison et al. 2007). Backtransforming estimates of net turnover and extinction fraction shows that HiSSE suffers the same precision issues in regards to the rate of extinction (Fig. 3, Fig. S2). Recently, it was reported that biases in the tip state ratios could also impact all parameter estimates (see Davis et al. 2013). Our simulations did not specifically test issues related to tip state biases. However, we did find that with HiSSE, the precision in extinction rate estimates remains fairly low regardless of the ratio of the states at the tips, especially when estimating extinction for the hidden state (Fig. 3). Interestingly, the lack of precision for extinction rates seems to have a relatively minor impact on estimates of net turnover or net diversification (Fig. 2-4; Fig. S1), though there is a general bias for net diversification to be underestimated as a consequence of inflated extinction rates. Nevertheless, it appears HiSSE correctly and qualitatively distinguishes differences in diversification among the various states in the model. And, as the number of species increases, the trajectory of the downward bias in net diversification suggests it will eventually disappear, and rate differences can be distinguished even if the differences are trivial (e.g., Fig. 2c).

For parameter estimates from the simulations testing both the CID models and the general HiSSE model we found very similar results as those described above, in that the model-averaged net turnover and net diversification parameters largely resembled the generating model (Fig. 4, S4). However, similar to the first set of simulations, there is a large amount of uncertainty and upward biases in estimates of extinction fraction. Thus, HiSSE generally has very low power in distinguishing multiple extinction fractions regardless of the number of states included in the model.

One issue of concern from our first set of simulations is that transition rates are almost always overestimated. This behavior appears unique to the HiSSE model in our simulations (Fig. 2), given that when evaluating data sets under BiSSE scenarios, transition rates are estimated reasonably well (Fig. S3). One suggestion from Rich FitzJohn (personal communication) is that this can occur when some states are present in low frequency, and since HiSSE has more states than BiSSE, it is likely that many state combinations are in very low frequencies. There are also relatively large confidence intervals surrounding each of the transition rate estimates that naturally favor models that assume equal transition rates, which should be reflected in the model-averaged rates. Indeed, as in the case of the HiSSE scenario that assumed pronounced differences in the transition rates, even 800 taxa was still not enough to unequivocally reject models that assumed equal transition rates (Table 1). We also examined the impact each parameter had on a fixed set of equilibrium frequencies, by randomly sampling sets of values and retaining those that estimated the same frequency within a small measure of error (see Supplemental Materials). The proportional range of values for the transition rates were more than double those found for the speciation rate, indicating that state frequencies are fairly resilient to changes in the transition rates. Taken together, estimating unique transition rates (and to some extent the rate of extinction) appears to be a difficult problem, which is not made any easier by HiSSE’s increase in state space. That is to say, HiSSE requires more from the data by including additional parameters without providing any more observable information. It is likely that in many cases far larger data sets with many more state origins than the ones we have generated here may be required to adequately estimate these particular parameters.

Finally, in regards to inference of ancestral states from the model-averaged parameter estimates, the simulations indicate that HiSSE correctly identifies and locates regions of the tree where supposed diversification rate differences have taken place (Fig. 5). The degree of reliability does, of course, depend on the size of the data set. For the HiSSE scenario that assumed a doubled speciation rate for state 1*B,* for example, data sets comprised of 50 taxa, 84.1% of the nodes, on average, have the state correctly inferred, and data sets comprised of 400 taxa 92.4% are correct. However, we note that there is also a general tendency for HiSSE to infer high marginal probabilities for the incorrect state (e.g., Fig. 5), which could provide misleadingly confident state reconstructions.

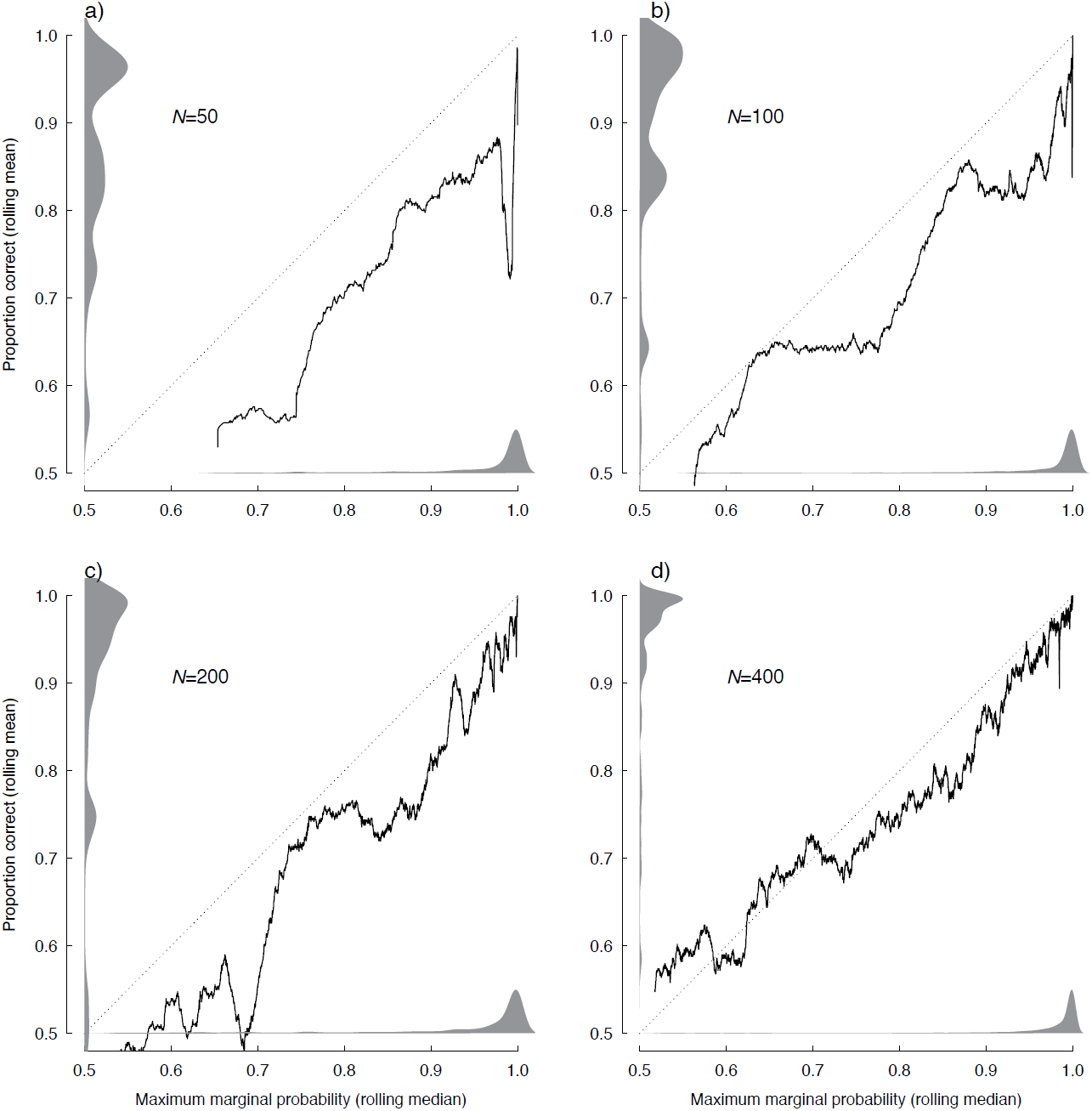

## Model rejection properties

We have strongly advocated (e.g., Beaulieu et al. 2012), as have many others (e.g., Anderson et al. 2000; Nickerson 2000; Nakagawa and Cuthill 2007), that parameter estimation is more important for understanding biology than is rejecting models. Model selection is part of this, of course, but it is more appropriate to examine a set of models and average or integrate their results, either through the use of information theory (e.g., Burnham and Anderson 2002), as we do here, or Bayesian approaches (e.g., Huelsenbeck et al. 2004). However, there are clearly still many studies whose main focus is rejecting trivial null models. The use of AIC or Akaike weights are inappropriate for these purposes (Burnham and Anderson 2002), despite being used frequently. Here, we examine how HiSSE and its various models perform in this way, especially in relation to the situation highlighted by Rabosky & Goldberg (2015), where a species tree evolves under a complex, unknown diversification process but traits evolve under a simple model independent of diversification parameters (e.g., our “worst-case” scenario).

Even though examining ΔAIC values are not recommended for model rejection, the rule of thumb from Burnham and Anderson (2002) remains popular for this use in phylogenetics: 0-2 means a model has substantial support, 4-7 means considerably less support, and >10 means essentially no support. Using these guidelines, we examined two of our simulations, the CID-2 and our “worst-case” scenario, both of which assume trait-independent diversification model (see Table S1). If we were to compare the fit of a constant birth-death BiSSE with a BiSSE model that assumes trait-dependent diversification, as Rabosky and Goldberg (2015) did, we found that when the true generating model was our CID-2 model, 38% of the time the BiSSE model had substantial support; in our “worst-case” scenario, 80% of the time BiSSE had substantial support. However, if we were to add just our CID-2 model into the set, in both cases the number of simulated data sets that show substantial support for the BiSSE case drop dramatically: just 7% in the CID-2 simulations, and just 1% in the “worst-case” scenario. If we examine the full set, which also includes CID-4 and the HiSSE model, 10% of simulated data sets that favor a trait-dependent scenario for CID-2, and 16% of the time under the “worst-case” scenario (Table S1).

We also conducted similar tests by simulating neutral characters on two empirical trees – the cetacean tree used by Rabosky and Goldberg (2015), and our own empirical tree (see below) – and found very similar behavior (see Table S2). More importantly, for parameter estimation, using a set of models and doing parameter estimation using a weighted average results in largely accurate inferences. For the 87-taxon cetacean tree, across simulated trees the model-averaged diversification rate in state 0 was within 10% of the rate for state 1 in 87% of the simulations (and we expect substantial uncertainty in these estimates); for Dipsidae, in 95% of the simulations the estimated diversification rate for state 0 was within 0.5% of the rate for state 1. In other words, even if a trait-dependent model were chosen, a careful biologist who looks at parameter estimates would be unlikely to find a rate difference of half a percent biologically significant.

In summary, we have gone from 94% of the time choosing the “wrong” model, to anywhere from 7-16% when the model set includes any number of our HiSSE models. Of course, if the goal is to simply find the best model, the rather marked improvement afforded by our HiSSE framework may remain somewhat unsatisfying. Future attention could, therefore, be paid to determining the appropriate ΔAIC that would constitute “substantial support” when using HiSSE, as opposed to simply using a ΔAIC<2 cutoff, as we have done here. Nevertheless, if the goal is to understand the potential biological implications provided by the parameters estimated from these models, even though the “best” model may at times be incorrect, taking into account model uncertainty when summarizing the parameter estimates will ameliorate spurious interpretations of trait-dependent diversification when it does not exist (e.g., Fig. 4, Fig. S5).

## The evolution of achene fruits

The development of this model was inspired by results from recent empirical work that applied BiSSE to understand the macroevolutionary consequences of evolving particular fruit types within a large flowering plant clade (i.e., campanulids; Beaulieu and Donoghue 2013). This study investigated whether diversification rates differences could explain why more than 80% of campanulid species exhibit fruits that are indehiscent (i.e., do not open mechanically), dry, and contain only a single seed. From a terminological standpoint, these fruits were broadly referred to as “achene” or “achene-like” to unify the various terms used to identify the same basic fruit character configuration (e.g., “cypselas” of Asteraceae – sunflowers and their relatives – or the single-seeded “mericarps” of Apiaceae – carrots and their relatives). According to the BiSSE model, the preponderance of achene fruits within campanulids can be explained by strong differences in diversification rates, with achene lineages having a rate that was roughly three times higher than non-achene lineages.

While these results are seemingly straightforward, they are complicated by the fact that the correlation between net diversification rates and the achene character state differed among the major campanulid lineages and was driven entirely by the inclusion of Asterales clade (Beaulieu and Donoghue 2013). Within Apiales and Dipsacales, the two remaining major achene-bearing clades, the diversification rate differences were not significant. However, in both these clades there were qualitative differences in the predicted direction, likely as a consequence of one or more shifts in diversification nested within one of the major achene-bearing clades. Together, these point to a more complex scenario for the interaction between achene fruits and diversification patterns that is not being adequately explained by BiSSE.

We illustrate an empirical application of HiSSE by examining the Dipsidae (Paracryphiales+Dipsacales; Tank and Donoghue 2010) portion of the achene data set of Beaulieu and Donoghue (2013). Specifically, we used HiSSE to locate and “paint” potential areas within Dipsidae that may be inflating the estimates of net diversification rates for achene lineages as a whole. We modified the original data set of Beaulieu and Donoghue (2013) in three important ways. First, we re-estimated the original molecular branch lengths using PAUP* (Swofford 2000), as opposed to relying on the branch lengths from the original RAxML (Stamatakis 2006) analysis, because PAUP* provides better optimization precision. Second, the molecular branch lengths were re-scaled in units of time using *treePL* (Smith and O’Meara 2012), an implementation of the penalized likelihood dating method of Sanderson (2002) specifically designed for large trees. We applied the same temporal constraints for Dipsidae as in the original Beaulieu and Donoghue (2013) study, and used cross-validation to determine the smoothing value that best predicted the rates of terminal branches pruned from the tree. Third, we conservatively removed various taxa of dubious taxonomic distinction, taxa considered varietals or subspecies of a species already contained within the tree, and tips that *treePL* assigned very short branch lengths (i.e., <1.0 Myr) – all of which have the tendency to negatively impact accuracy of estimating diversification rates (though in this case, rerunning the analyses including such tips did not have a qualitative effect on the results). The exclusion of these taxa resulted in a data set comprised of 417 species from the original 457.

We fit 24 different models to the achene data set for Dipsidae. Four of these models corresponded to BiSSE models that either removed or constrained particular parameters, 16 corresponded to various HiSSE models that assumed a hidden state associated with both the observed states (i.e., non-achene+, achene+), and four corresponded to various forms of our trait-independent models (i.e., CID-2, CID-4). In all cases, we incorporated a unique sampling frequency scheme to the model. Rather than assuming random sampling for the entire tree (see above), we included the sampling frequency for each major clade included in the tree (see Table S3; note, that the results are qualitatively similar regardless of sampling scheme used). In order to generate a measure of confidence for parameters estimated under a given model, we implemented an “adaptive search” procedure that provides an estimate of the parameter space that is some pre-defined likelihood distance (e.g., 2 lnL units) from the maximum likelihood estimate (MLE), which follows from Edwards (1992). We also took into account both state uncertainty and uncertainty in the model when “painting” diversification rates across the tree. Our procedure first calculated a weighted average of the likeliest state and rate combination for every node and tip for each model in the set, using the marginal probability as the weights, which were then averaged across all models using the Akaike weights. All analyses were carried out in *hisse.*

The best model, based on Akaike weights, was a relatively simple HiSSE model with regards to the number of free parameters it contained (Table 3). The model suggests character-dependent diversification with fruit type, where only rates for non-achene+ and achene+ are allowed to be free, and where transitions between state 0*B* and 1*B* were disallowed. This model had a pronounced improvement over the set of BiSSE models, where none had an Akaike weight that was greater than 0.001. However, before describing the parameter estimates of the best model, we note that, with a modified tree and character set, the parameter estimates under the BiSSE models were different from those reported by Beaulieu and Donoghue (2013). Specifically, the higher diversification rates estimated for achene lineages (*r*_achene_=0.148, support range: [0.121,0.161]) compared to non-achene lineages (*r*_non-achene_=0.065, support range: [0.0498,0.083]) were indeed significant based on a sampling of points falling within 2lnL units away from the MLE. The parameters estimated under the HiSSE model, on the other hand, suggest a more nuanced interpretation of this result. The higher diversification rates of clades bearing achenes as a whole is likely due to a hidden state nested within some of these lineages that is associated with exceptionally high diversification rates (*r*_achene+_=0.199, support region: [0.179,0.221]). In fact, the model suggests that achene lineages not associated with the high diversification hidden state have a diversification rate that is indistinguishable from the non-achene diversification rate (*r*_non-achene−_= *r*_achene−_=0.059, support range: [0.049,0.068]). Non-achene lineages associated with the high diversification hidden state also show elevated diversification rates (*r*_non-achene+_=0.158, support region: [0.138,0.185]), relative to the rate of the non-achene state with the other hidden state, suggesting strong rate heterogeneity even in Dipsidae lineages that bear other fruit types.

**Table 3.** The fit of alternative models of achene fruit evolution in the flowering plant clade Dipsidae. The best model, based on ΔAIC and Akaike weights (*w_i_*) is denoted in bold.

| Model | np | lnLik | AIC | $\Delta AIC$ | $w_i$ |
| --- | --- | --- | --- | --- | --- |
| BiSSE: All free | 6 | -1486.5 | 2984.9 | 90.3 | <0.001 |
| BiSSE: $\epsilon_0 = \epsilon_1$ | 5 | -1488.6 | 2987.2 | 92.6 | <0.001 |
| BiSSE: q's equal | 5 | -1487.3 | 2984.5 | 89.9 | <0.001 |
| BiSSE: $\epsilon_0 = \epsilon_1$, q's equal | 4 | -1490.2 | 2988.4 | 93.8 | <0.001 |
| Null-two: q's equal | 5 | -1449.1 | 2908.3 | 13.7 | 0.001 |
| Null-two: $\epsilon$'s, q's equal | 4 | -1449.7 | 2907.4 | 12.8 | 0.001 |
| Null-four: q's equal | 9 | -1445.1 | 2908.1 | 13.5 | 0.001 |
| Null-four: $\epsilon$'s equal, q's equal | 6 | -1445.3 | 2902.6 | 7.98 | 0.015 |
| HiSSE: q's equal | 9 | -1446.6 | 2911.2 | 16.6 | <0.001 |
| HiSSE: $\epsilon$'s equal, q's equal | 6 | -1447.2 | 2906.5 | 11.9 | 0.002 |
| HiSSE: $\tau_0 A = \tau_1 A = \tau_0 B$, $\epsilon_0 A = \epsilon_1 A = \epsilon_0 B$, q's equal | 5 | -1463.3 | 2936.6 | 42.0 | <0.001 |
| HiSSE: $\tau_0 A = \tau_1 A = \tau_0 B$, $\epsilon$'s equal, q's equal | 4 | -1471.0 | 2950.0 | 55.4 | <0.001 |
| HiSSE: $q_0 B_1 B = 0$, $q_1 B_0 B = 0$, all other q's equal | 9 | -1451.7 | 2921.5 | 26.9 | <0.001 |
| HiSSE: $\epsilon$'s equal, $q_0 B_1 B = 0$, $q_1 B_0 B = 0$, all other q's equal | 6 | -1451.7 | 2915.5 | 20.9 | <0.001 |
| HiSSE: $\tau_0 A = \tau_1 A = \tau_0 B$, $\epsilon_0 A = \epsilon_1 A = \epsilon_0 B$, $q_0 B_1 B = 0$, $q_1 B_0 B = 0$, all other q's equal | 5 | -1462.2 | 2934.3 | 39.7 | <0.001 |
| HiSSE: $\tau_0 A = \tau_1 A = \tau_0 B$, $\epsilon$'s equal, $q_0 B_1 B = 0$, $q_1 B_0 B = 0$, all other q's equal | 4 | -1469.5 | 2947.1 | 52.5 | <0.001 |
| HiSSE: $\tau_0 A = \tau_0 B$, $\epsilon_0 A = \epsilon_0 B$, q's equal | 7 | -1459.6 | 2933.2 | 38.6 | <0.001 |
| HiSSE: $\tau_0 A = \tau_0 B$, $\epsilon$'s equal, q's equal | 5 | -1466.4 | 2942.8 | 48.2 | <0.001 |
| HiSSE: $\tau_0 A = \tau_0 B$, $\epsilon_0 A = \epsilon_0 B$, $q_0 B_1 B = 0$, $q_1 B_0 B = 0$, all other q's equal | 7 | -1458.4 | 2930.8 | 36.2 | <0.001 |
| HiSSE: $\tau_0 A = \tau_0 B$, $\epsilon$'s equal, $q_0 B_1 B = 0$, $q_1 B_0 B = 0$, all other q's equal | 5 | -1464.7 | 2939.4 | 44.8 | <0.001 |
| HiSSE: $\tau_0 A = \tau_1 A$, $\epsilon_0 A = \epsilon_1 A$, q's equal | 7 | -1446.7 | 2907.3 | 12.7 | <0.001 |
| HiSSE: $\tau_0 A = \tau_1 A$, $\epsilon$'s equal, q's equal | 5 | -1469.1 | 2948.3 | 53.7 | <0.001 |
| HiSSE: $\tau_0 A = \tau_1 A$, $\epsilon_0 A = \epsilon_1 A$, $q_0 B_1 B = 0$, $q_1 B_0 B = 0$, all other q's equal | 7 | -1442.2 | 2898.3 | 3.71 | 0.130 |
| HiSSE: $\tau_0 A = \tau_1 A$, $\epsilon$'s equal, $q_0 B_1 B = 0$, $q_1 B_0 B = 0$, all other q's equal | 5 | **-1442.3** | **2894.6** | **0.00** | **0.848** |

Character reconstructions identified two transitions to the hidden fast state in achene lineages, and thus a higher diversification rate: one shift occurred along the stem leading to crown Dipsacaceae and the other occurred along the stem leading to “core Valerianaceae” (the most inclusive clade that excludes *Patrinia, Nardostachys,* and *Valeriana celtica)* (Fig. 6). It is important to note that in the case of non-achene lineages, the model identified four shifts to the hidden fast state – one at the base of Caprifolieae *(Lonicera, Symphoricarpus, Leycesteria,* and *Triosteum),* and three within *Viburnum* (Fig. 6). The several shifts detected in *Viburnum* are noteworthy in that they correspond almost exactly with the inferred shifts detected in a more focused study of the genus that applied various models of diversification (Spriggs et al. 2015).

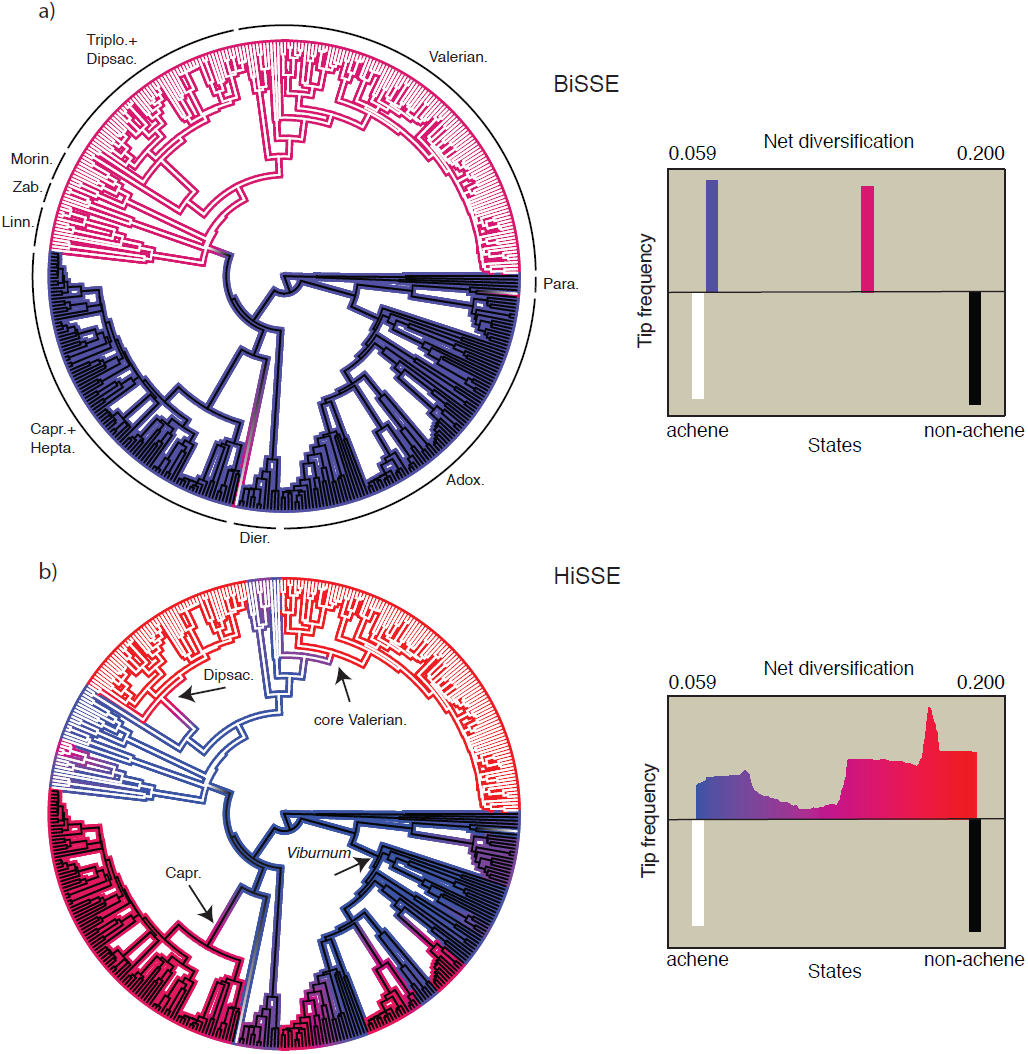

In regards to achene lineages, the general location of the inferred shifts is intriguing, as they appear to coincide with specific clades that exhibit specialized structures related to achene dispersal. In Dipsacaceae, for example, there is tremendous diversity in the shape of the “epicalyx”, a tubular structure that subtends the calyx and encloses the ovary (Donoghue et al. 2003; Carlson et al. 2009), which is often associated with elaborated structures (e.g., “wings”, “pappus-like” bristles) that accompany their achene fruits. Interestingly, many of these same general forms are observed throughout core Valerianaceae, although they arise from modifications to the calyx (Donoghue et al. 2003; Jacobs et al. 2010). While the significance of these structures is thought to improve protection, germination, and dispersal of the seed, we emphasize that, at this stage, it is difficult to confidently rule out other important factors, such important biogeographic movements due to increased dispersal abilities (i.e., “key opportunities” *sensu* Moore and Donoghue 2007), or even purely genetic changes, such as gene and genome duplications (Hildago et al. 2010; Carlson et al. 2011). But, we can at least confidently conclude that the achene by itself is likely not a strong correlate of diversity patterns within Dipsidae.

## Discussion

Progress in biology comes from confronting reality with our hypotheses and either confirming that our view of the world is correct, or, more excitingly, finding out that we still have a lot left to discover. With studies of diversification, we have mostly been limited to doing the former – we have an idea about a trait that may affect diversification rates based on intuition about its potential effect (e.g., an achene might allow greater dispersal, and thus easier colonization of new habitats to form new species) as well as some knowledge of its distribution (e.g., some very large plant clades have achenes) and then run an analysis. Typical outcomes are yes, there is a difference in diversification in the way we expected, or there is no difference but maybe we just lack the necessary power. In any case, chances are we are at least vaguely correct that the trait we think credibly has a mechanism for increasing diversification rate actually has an effect (at least once there is enough power), or, less compellingly, clades we identify for such a test from eyeballing the data return a significant result despite no underlying reality.

Surprise is a necessary part of discovery that, to put it bluntly, has been relatively lacking in trait-dependent diversification studies until now. With HiSSE we can still test our intuitions about a particular character, but we can also discover that rates seem to be driven by some unknown and unmeasured character state, allowing the data to help us generate new hypotheses – Is the diversification rate correlated with the achene, or is the achene simply a necessary precursor to some other trait that is more likely to be driving diversification? This lets us go from a scenario where we simply reject trivial nulls, such as whether diversification rates of clades with and without some focal trait are precisely equal, to being potentially surprised by the results – No, it is not the achene *per se,* but it is something else nested within these particular clades in addition to the achene fruit type.

Currently, diversification models are divided between those that look at one or more focal traits only, integrating over any other factors (e.g., BiSSE, Maddison et al. 2007; BiSSE-ness, Magnuson-Ford and Otto, 2012; ClaSSE, Goldberg and Igic, 2012; sister group comparisons, Mitter et al. 1988), and those that fit rates to trees but ignore trait information altogether (e.g., MEDUSA, Alfaro et al. 2009; BAMM, Rabosky 2014). Our HiSSE framework spans this range. If the true model is strictly trait-dependent diversification, it can detect this (Fig. 1-5), as a BiSSE analysis would. If the true model is rates varying due to some other unexamined factor (a physical trait, a property of the environment, etc.) HiSSE recovers this too, as in the case of achene evolution in the Dipsidae clade (Fig. 6). Uniquely, HiSSE can also give an intermediate answer where a focal trait explains some but not all of the diversification difference.

The HiSSE framework also addresses some of the recent important criticisms levied against state speciation and extinction models (Rabosky and Goldberg 2015). Specifically, our method no longer requires the assumption that focal character states are associated with diversification rate differences. Instead, it allows this assumption to be explored as part of a more flexible overall model, as opposed to relying on separate tests for uncovering character-dependent and CID rates of diversification (e.g., Beaulieu and Donoghue 2013; Weber and Agrawal 2014; Spriggs et al. 2015). Including our models of trait-independent diversification in a set of alternative trait-dependent models should also alleviate concerns of spurious assignments of diversification rate differences between observed character states in cases where trees are evolving separately from the focal trait (c.f., Rabosky and Goldberg 2015). These CID diversification models are designed specifically to be as complex as competing BiSSE or HiSSE models, which provides a fairer comparison over more trivial equal-rate “nulls.”

We do, however, highlight one important statistical concern in which HiSSE requires significant caution in its use. This involves its indifference to number of changes (c.f., Maddison and FitzJohn 2014). As with BiSSE, we should find it more credible that a particular character state enhances diversification rates if we see a rate increase in each of the 10 times it evolves than if it evolved just once but had the same magnitude of rate increase. A good solution to this problem has not yet been proposed, and we urge a healthy skepticism of any result based on a trait that has evolved only a few times.

There are other practical concerns in empirical applications of the HiSSE model. While it could be used over a set of trees (i.e., bootstrap or Bayesian post-burnin tree samples), the model assumes that the branch lengths, topology, and states are known without error. Certain kinds of phylogenetic errors, such as terminal branch lengths that are too long (as may occur with sequencing errors in the data used to make the tree) can result in particular biases in estimates of speciation and/or extinction (also see Beaulieu and O’Meara 2015). Similarly, if one clade were reconstructed to be younger than it actually is, due to a substitution rate slowdown caused by some other trait (e.g., life-history; Smith and Donoghue 2008), it could be interpreted as having a faster diversification rate, perhaps even inferred to have its own hidden state. For trees that come from Bayesian dating analyses, whether a birth-death or Yule prior was used may affect results unless the data is strong enough to overwhelm the prior (Condamine et al. 2015). Future simulation studies will focus on better understanding the impact that these and other branch length error scenarios can have on the interpretation and estimation of various model parameters. Also, we point out that the HiSSE model assumes discrete characters, whether they be hidden or observed, but it could be that a continuous parameter is the cause of a diversification rate difference (e.g., perhaps extinction risk varies inversely with mean of the dispersal kernel). However, this may only enter the model as an unseen discrete character, perhaps corresponding to low and high values of the continuous character.

Our HiSSE model is part of a long tradition in comparative methods of identifying, and then addressing, perceived shortcomings. For instance, Felsenstein (1985) pointed out the pitfalls of treating species values as independent data points and provided two alternatives (i.e., independent contrasts and dividing a tree into pairs of tips) to address those issues. Maddison (2006) realized the issue of not accounting for diversification in transition-based methods (i.e., Pagel 1994) and not accounting for differential transitions in diversification models, which led to the development of state-dependent speciation and extinction models (Maddison et al. 2007; FitzJohn et al. 2009). Recently, Rabosky and Goldberg (2015) pointed out a serious problem with interpretations in state speciation and extinction analyses in general, and our work largely, but not entirely, addresses these concerns. We caution users that it is important to keep a perspective: all methods have flaws, and all will fail given a strong enough violation of their underlying assumptions. For example, it appears difficult for HiSSE to adequately estimate different transition rates when the model assumes any number of hidden states, and so many estimates will be biased. Even with the increasing efforts to test new methods (in our case, over 17,000 computer-days were devoted to conducting simulations and analyses) there will be flaws that may have gone undetected. We urge skepticism towards all models, but also skepticism towards statements of fatal flaws in some models while leaving newer, competing models relatively untested.

There is no question that state-dependent speciation and extinction models are an important advancement for understanding characters’ impact on diversification patterns. They have greatly improved statistical power over older, simpler sister-clade comparisons, and the explicit inference of differences in speciation and extinction has the potential for a much more fine-grained analysis of diversification. But, in a way, these models have also allowed us to retreat to the old comforts of reducing complex organisms into units of single, independently evolving characters, and offering adaptive interpretations to each (c.f., Gould and Lewontin 1979). To be fair, of course, it is unlikely that any trait of great interest to biologists has exactly zero effect on speciation and/or extinction rates, but it is certainly unlikely that this trait acts in isolation. Thus, we hope HiSSE is viewed as a step away from this line of thinking, as we no longer have to necessarily focus analyses, or even interpret the results, by reference to the focal trait by itself, but can instead estimate how important it is as a component of diversification overall. It is in this way that analyses focused on “hidden” factors promoting diversification will afford us a more refined understanding of why certain clades become extraordinarily diverse, while still allowing us to examine our hypotheses about effects of observed traits.

## Supplemental Material

Data available from the Dryad Digital Repository: http://dx.doi.org/10.5061/dryad.52hp1

## Funding

This work was supported by the National Institute for Mathematical and Biological Synthesis [to J.M.B.], an Institute sponsored by the National Science Foundation, the U.S. Department of Homeland Security, and the U.S. Department of Agriculture through NSF Award #EF-0832858, with additional support from The University of Tennessee, Knoxville.

## Acknowledgements

We thank Nathan Jackson, Nick Matzke, Kathryn Massana, Chuck Bell, Andrew Leslie, Jim Fordyce, and Chris Hamm for helpful discussions at various stages of this manuscript. Rich FitzJohn, Sally Otto, and an anonymous reviewer provided very useful comments on the manuscript, and we are deeply indebted to Tanja Stadler for her suggestion to consider more complex null models, which led to the CID models here. We also thank Jeffrey Oliver and Elizabeth Spriggs for feedback on the method implementation. JMB would like to thank Michael Donoghue for wonderful discussions over the years regarding some of the concepts described here.

## References

Alexander H.K., Lambert A., Stadler T. 2015. Quantifying age-dependent extinction from species phylogenies. Systematic Biology In press. doi:10.1093/sysbio/syv065.

Alfaro M.E., Santini F., Brock C., Alamillo H., Dornbug A., Rabosky D.L., Carnevale G., Harmon L.J. 2009. Nine exceptional radiations plus high turnover explain species diversity in jawed vertebrates. Proceedings of the National Academy of Sciences of the U.S.A. 106: 13410–13414.

Anderson D.R., Burnham K.P., Thompson W.L. 2000. Null hypothesis testing: problems, prevalence, and an alternative. Journal of Wildlife Management 64: 912–923.

Beaulieu J.M., Donoghue M.J. 2013. Fruit evolution and diversification in campanulid angiosperms. Evolution 67: 3132–3144.

Beaulieu J.M., Jhueng D.-C., Boettiger C., O’Meara B.C. 2012. Modeling stabilizing selection: expanding the Orstein-Uhlenbeck model of adaptive evolution. Evolution 66: 2369–2383.

Beaulieu J.M., O’Meara B.C., Donoghue M.J. 2013. Identifying hidden rate changes in the evolution of a binary morphological character: the evolution of plant habit in campanulid angiosperms. Systematic Biology 62: 725–737.

Beaulieu J.M., O’Meara B.C. 2014. Hidden Markov models for studying the evolution of binary morphological characters. In Modern Phylogenetic Comparative Methods and their Application in Evolutionary Biology, Garamszegi L.Z. editor. London:Springer.

Beaulieu J.M., O’Meara B.C. 2015. Extinction can be estimated from moderately sized molecular phylogenies. Evolution 69: 1036–1043.

Burnham K.P., Anderson D.R. 2002. Model selection and multimodel inference: a practical information-theoretic approach. New York:Springer.

Carlson S.E., Mayer V., Donoghue M.J. 2009. Phylogenetic relationships, taxonomy, and morphological evolution in Dipsacaceae (Dipsacales) inferred by DNA sequence data. Taxon 58: 1075–1091.

Carlson S.E., Howarth D.G., Donoghue M.J. 2011. Diversification of CYCLOIDEA-like genes in Dipscaceae (Dipsacales): implications for the evolution of capitulum inflorescences. BMC Evolutionary Biology 11: 325.

Condamine F.L., Nagalingum N.S., Marshall C.R., Morlon H. 2015. Origin and diversification of living cycads: a cautionary tale on the impact of the branching process prior in Bayesian molecular dating. BMC Evolutionary Biology 15: 65.

Davis M.P., Midford P.E., Maddison W. 2013. Exploring power and parameter estimation of the BiSSE method for analyzing species diversification. BMC Evolutionary Biology 13: 38.

de Querioz A. 2002. Contingent predictability in evolution: Key traits and diversification. Systematic Biology 51: 917–929.

Donoghue M.J., Bell C.D., Winkworth R.C. 2003. The evolution of reproductive characters in Dipsacales. International Journal of Plant Sciences. 164:S453–S464.

Donoghue M.J., Sanderson M.J. 2015. Confluence, synnovation, and depauperons in plant diversification. New Phytologist doi:10.1111/nph.13367.

Edwards A.W.F. 1992. Likelihood. Cambridge:Cambridge University Press.

Felsenstein J. 1985. Phylogenies and the comparative method. The American Naturalist 125: 1–15.

FitzJohn R.G. 2012. Diversitree: comparative phylogenetic analyses of diversification in R. Methods in Ecology and Evolution 3: 1084–1092.

FitzJohn R.G., Maddison W.P., Otto S.P. 2009. Estimating trait-dependent speciation and extinction rates from incompletely resolved phylogenies. Systematic Biology 58: 595–611.

Goldberg E.E., Kohn J.R., Lande R., Robertson K.A., Smith S.A., Igic B. 2010. Species selection maintains self-incompatibility. Science 330: 493–495.

Goldberg E. E., Igic B. 2012. Tempo and mode in plant breeding system evolution. Evolution 66: 3701–3709.

Gould S.J., Lewontin R.C. 1979. The spandrels of San Marco and the Panglossian paradigm: a critique of the adaptationist programme. Proceedings of the Royal Society, B. 205: 581–598.

Hansen T.F. 1997. Stabilizing selection and the comparative analysis of adaptation. Evolution 51: 1341–1351.

Hardy N.B. and Otto S.P. 2014. Specialization and generalization in the diversification of phytophagous insects: tests of the musical chairs and oscillation hypotheses. Proceedings of the Royal Society B. 281: 20132960.

Harmon L.J., Weir J.T., Brock C.D., Glor R.E., Challenger W. 2008. GEIGER: investigating evolutionary radiations. Bioinformatics 24: 129–131.

Hildago O., Mathez J., Garcia S., Garnatje T., Pellicer J., Valles J. 2010. Genome size study in the Valerianaceae: First results and new hypotheses. Journal of Botany. doi:10.1155/2010/797246.

Huelsenbeck J.P., Larget B., Alfaro M.E. 2004. Bayesian phylogenetic model selection using reversible jump Markov chain Monte Carlo. Molecular Biology and Evolution 21: 1123–1133.

Jacobs B., Bell C., Smets E. 2010. Fruits and seeds of the Valeriana clade (Dipsacales): Diversity and evolution. International Journal of Plant Sciences 171: 421–434.

Maddison, W. P. 2006. Confounding asymmetries in evolutionary diversification and character change. Evolution 60: 1743–1746.

Maddison W.P., Midford P.E., Otto S.P. 2007. Estimating a binary character’s effect on speciation and extinction. Systematic Biology 56: 701–710.

Maddison W.P., FitzJohn R.G. 2014. The unsolved challenge to phylogenetic correlation tests for categorical characters. Systematic Biology 64: 127–136.

Magnuson-Ford K., Otto S.P. Linking the investigations of character evolution and species diversification. American Naturalist 180: 225–245.

Marazzi B., Ane C., Simon M.F., Delgado-Salinas A., Luckow M., Sanderson M.J. 2012. Locating evolutionary precursors on a phylogenetic tree. Evolution 66: 3918–3930.

Mitter C., Farrell B., Wiegmann B. 1988. The phylogenetic study of adaptive zones: Has phytophagy promoted insect diversification? American Naturalist 132: 107–128.

Moore B.R., Donoghue M.J. 2007. Correlates of diversification in the plant clade Dipsacales: geographic movement and evolutionary innovations. American Naturalist 170:S28–S55.

Morlon H., Parsons T.L., Plotkin J.B. 2011. Reconciling molecular phylogenies with the fossil record. Proceedings of the National Academy of Sciences of the U.S.A. 108: 16327–16332.

Nakagawa S., Cuthill I.C. 2007. Effect size, confidence interval and statistical significance: a practical guide for biologists. Biological Reviews 82: 591–605.

Nee S., Holmes E.C., May R.M., Harvey P.H. 1994. Extinction rates can be estimated from molecular phylogenies. Philosophical Transactions of the Royal Society, B. 344: 77–82.

Nickerson R.S. 2000. Null hypothesis significance testing: a review of an old and continuing controversy. Psychological Methods 5: 241–301.

Pagel M. 1994. Detecting correlated evolution on phylogenies: a general method for the comparative analysis of discrete characters. Proceedings of the Royal Society, B. 255: 37–45.

Pennell M.W., Eastman J.M., Slater G.J., Brown J.W., Uyeda J.C., FitzJohn R.G., Alfaro M.E., Harmon L.J. 2014. Geiger v2.0: an expanded suite of methods for fitting macroevolutionary models to phylogenetic trees. Bioinformatics doi:10.1093/bioinformatics/btu181

Price S.A., Hopkins S.S.B, Smith K.K., Roth V.L. 2012. Tempo of trophic evolution and its impact on mammalian diversification. Proceedings of the National Academy of Sciences of the U.S.A. 109: 7008–7012.

Rabosky D.L. 2010. Extinction rates should not be estimated from molecular phylogenies. Evolution 64: 1816–1824.

Rabosky D.L. 2014. Automatic detection of key innovations, rate shifts, and diversity-dependence on phylogenetic trees. PLoS One doi:10.1371/journal.pone.0089543

Rabosky D.L., Lovette I.J. 2008. Explosive evolutionary radiations: decreasing speciation or increasing extinction through time? Evolution 62: 1866–1875.

Rabosky D.L., Goldberg E.E. 2015. Model inadequacy and mistaken inferences of trait-dependent speciation. Systematic Biology 64: 340–355.

Sanderson M.J. 2002. Estimating absolute rates of molecular evolution and divergence times: a penalized likelihood approach. Molecular Biology and Evolution 19: 101–109.

Schluter D., Price T., Mooers A.O., Ludwig D. 1997. Likelihood of ancestor states in adaptive radiation. Evolution 51: 1699–1711.

Slater G.J., Price S.A., Santini F., Alfaro M.E. 2010. Diversity versus disparity and the radiation of modern cetaceans. Proceedings of the Royal Society, B. 277: 3097–3104.

Smith S. A., Donoghue M.J. 2008. Rates of molecular evolution are linked to life history in flowering plants. Science 322: 86–89.

Smith S.A. O’Meara B.C. 2012. treePL: Divergence time estimation using penalized likelihood for large phylogenies. Bioinformatics 28: 2689–2690.

Spriggs E.L., Clement W.L., Sweeney P.W., Madriñán, Edwards E.J., Donoghue M.J. 2015. Temperate radiations and dying embersof a tropical past: the diversification of Viburnum. New Phytologist. In press, doi:10.1111/nph.13305.

Stamatakis A. 2006. Maximum likelihood-based phylogenetic analyses with thousands of taxa and mixed models. Bioinformatics 22: 2688–2690.

Swofford D.L. 2000. PAUP*: Phylogenetic analysis using parsimony (and other methods) version 4. Sinauer Associates, Sunderland, Massachusetts.

Tank D.C., Donoghue M.J. 2010. Phylogeny and phylogenetic nomenclature of the Campanulidae based on an expanded sample of genes and taxa. Systematic Botany 35: 425–441.

Weber M.G., Agrawal A.A. 2014. Defense mutualisms enhance plant diversification. Proceedings of the National Academy of Sciences of the U.S.A. 111: 1644216447.

Wilson A.W., Binder M., Hibbett D.S. 2011. Effects of gasteroid fruiting body morphology on diversification rates in three independent clades of fungi estimated using binary state speciation and extinction analysis. Evolution 65: 1305–1322.

Yang Z., Kumar S., Nei M. 1995. A new method of inference of ancestral nucleotide and amino acid sequences. Genetics 141: 1641–1650.

